# Dense cellular segmentation for EM using 2D-3D neural network ensembles

**DOI:** 10.1101/2020.01.05.895003

**Authors:** Matthew D. Guay, Zeyad A.S. Emam, Adam B. Anderson, Maria A. Aronova, Brian Storrie, Irina D. Pokrovskaya, Richard D. Leapman

## Abstract

Cell biologists can now build 3D models from segmentations of electron microscopy (EM) images, but accurate manual segmentation of densely-packed organelles across gigavoxel image volumes is infeasible. Here, we introduce 2D-3D neural network ensembles that produce dense cellular segmentations at scale, with accuracy levels that outperform baseline methods and approach those of human annotators.

---

Biomedical researchers use electron microscopy (EM) to image cells, organelles, and their constituents at the nanoscale. The serial block-face scanning electron microscope (SBF-SEM)^1^ employs automated serial sectioning techniques on block samples to produce gigavoxel image volumes. This rapid growth in throughput challenges traditional image analytic workflows for EM, which rely on trained humans to identify salient image features. This paper develops the *dense cellular segmentation* method, which classifies each voxel in an image into categories from a detailed schema of cellular and subcellular structures. Structural biologists can use dense cellular segmentations to provide rich 3D ultrastructural models yielding new insights into cellular processe^2,3^, but applying this method across entire SBF-SEM datasets is infeasible unless analytic bottlenecks are surmounted.

It is challenging to automate dense segmentation tasks for EM due to the image complexity of biological structures at the nanoscale. An image with little noise and high contrast between features may be accurately segmented with simple thresholding methods, while accurate segmentation of images with multiscale features, noise, and textural content remains an open problem for many biomedical applications. The need for solutions to such problems has driven seminal contributions to image segmentation literature from EM microscopy, including the U-Net^4^, 3D U-Net variants^5, 6^, and newer volumetric segmentation algorithms^7-11^, as well as recent plug-and-play tools such as CDeep3M^12^. We introduce a new 3D biomedical segmentation algorithm based on ensembles of neural networks with separated 2D and 3D convolutional modules and prediction heads, building off of existing work in volumetric image segmentation^9, 13, 14^ to solve a dense segmentation task for SBF-SEM datasets of human blood platelets, as a model system. We show that our algorithm outperforms baselines, has comparable image quality to our human annotators, and closely matches human performance on a downstream biological analysis task. We use the algorithm to segment a billion-voxel block sample in an hour on a single NVIDIA GTX 1080 GPU, demonstrating a segmentation capability that is infeasible without automation and accessible to commodity computing tools.

The datasets used in this study are 3D electron microscopy image volumes from two human platelet samples: Subject1 and Subject2. We focus on four data subsets: *training data* (50 × 800 × 800 voxels), *evaluation data* (24 × 800 × 800 voxels), *test data* (121 × 609 × 400 voxels), and *annotator comparison data* (110 × 602 × 509 voxels). The training and evaluation data are subvolumes taken from Subject1. The test data and annotator comparison data are taken from Subject2. The manually-annotated ground truth labels for the training and evaluation datasets cover the entirety of their respective image volumes, while the test and annotator comparison labels each cover a single cell within the image. The labeling schema divides image content into seven classes, each with an assigned color: `background` (black), `cell` (darkgreen), `mitochondrion` (magenta), `canalicular channel` (yellow), `alpha granule` (dark blue), `dense granule` (bright red), and `dense granule core` (dark red). Figure S1 shows sample images of the datasets and ground truth labels.

Inspired by existing work on combining 2D and 3D computations for volumetric data analysi^s13,14^ we experiment with combinations of 2D and 3D neural modules to trade off between computational efficiency and spatial context. The highest-performing network architecture in this paper, 2D-3D+3×3×3, is a composition of a 2D U-Net-style encoder-decoder and 3D convolutional spatial pyramid, with additional 3×3×3 convolutions at the beginning of convolution blocks in the encoder-decoder. We use same-padded convolution operations throughout, so that 3D operations can be used on anisotropic data windows with small size along the **z** axis.

Our best algorithms are ensembles that average the per-voxel class probability distributions across several networks. The ensembled networks are identical architectures trained from different random weight initializations. When describing segmentation algorithms, we use Top-**k** to indicate an ensemble of the best *k* instances of an architecture. Figure 2 details our best network architecture and illustrates the ensembling process.

**Figure 1.**
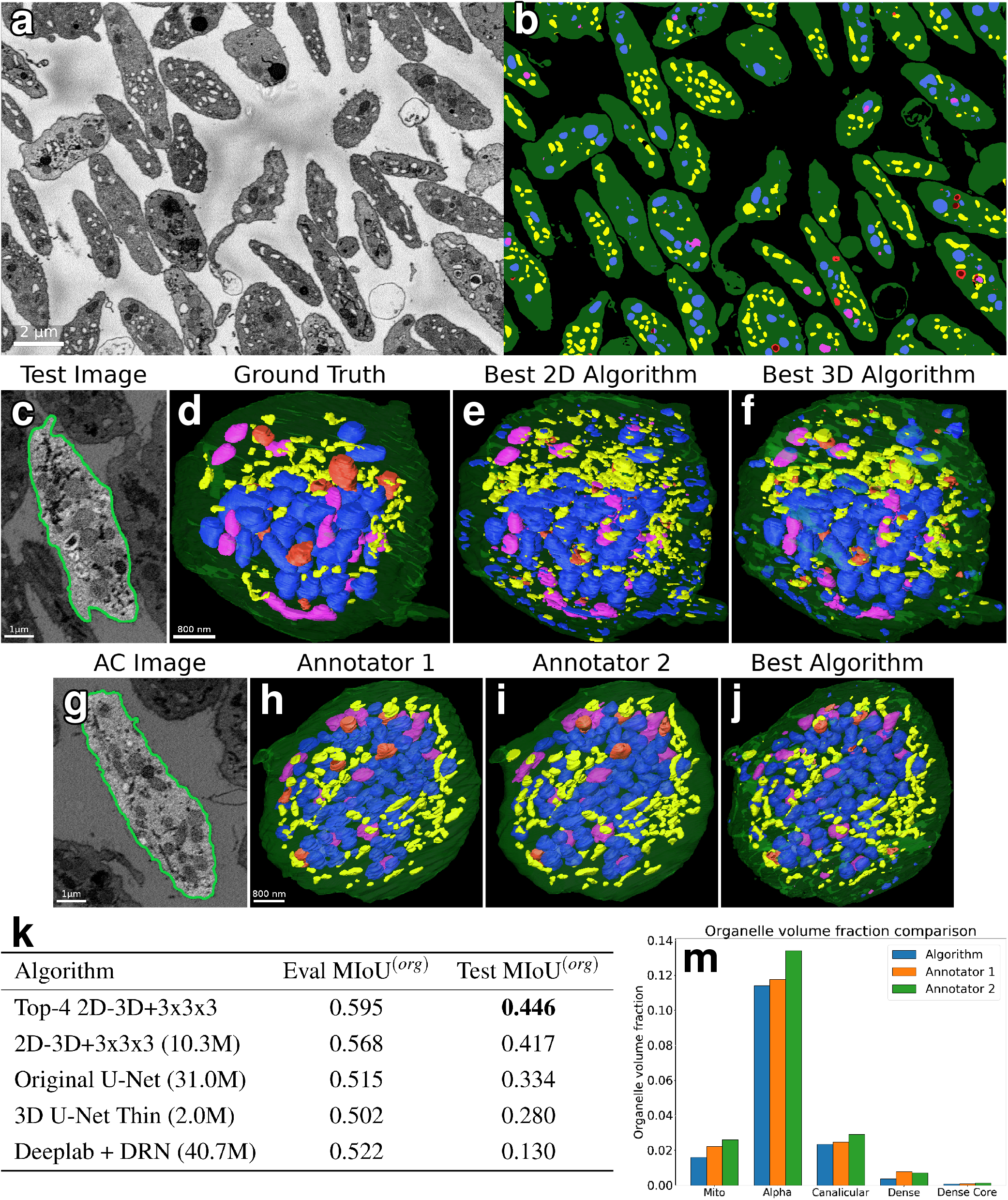
Results. (**a-b**) Orthoslice of Subject 1 image and segmentation. (**c**) Test dataset orthoslice, segmented cell highlighted. (**d-f**) Comparison between ground truth segmentation of test cell and our best 2D and 3D algorithms. (**g**) Annotator comparison (AC) dataset orthoslice, segmented cell highlighted. (**h-j**) Annotator comparison cell segmentations, comparing the two human annotators and our best algorithm. (**k**) Summarized comparison of mean intersection-over-union across organelle classes (MIoU^(*org*)^) on test and evaluation datasets for segmentation algorithms. For full results, see S1. (**m**) Comparison of organelle volume fractions between two human annotators and our best algorithm, computed from annotator comparison cell segmentations.

**Figure 2.**
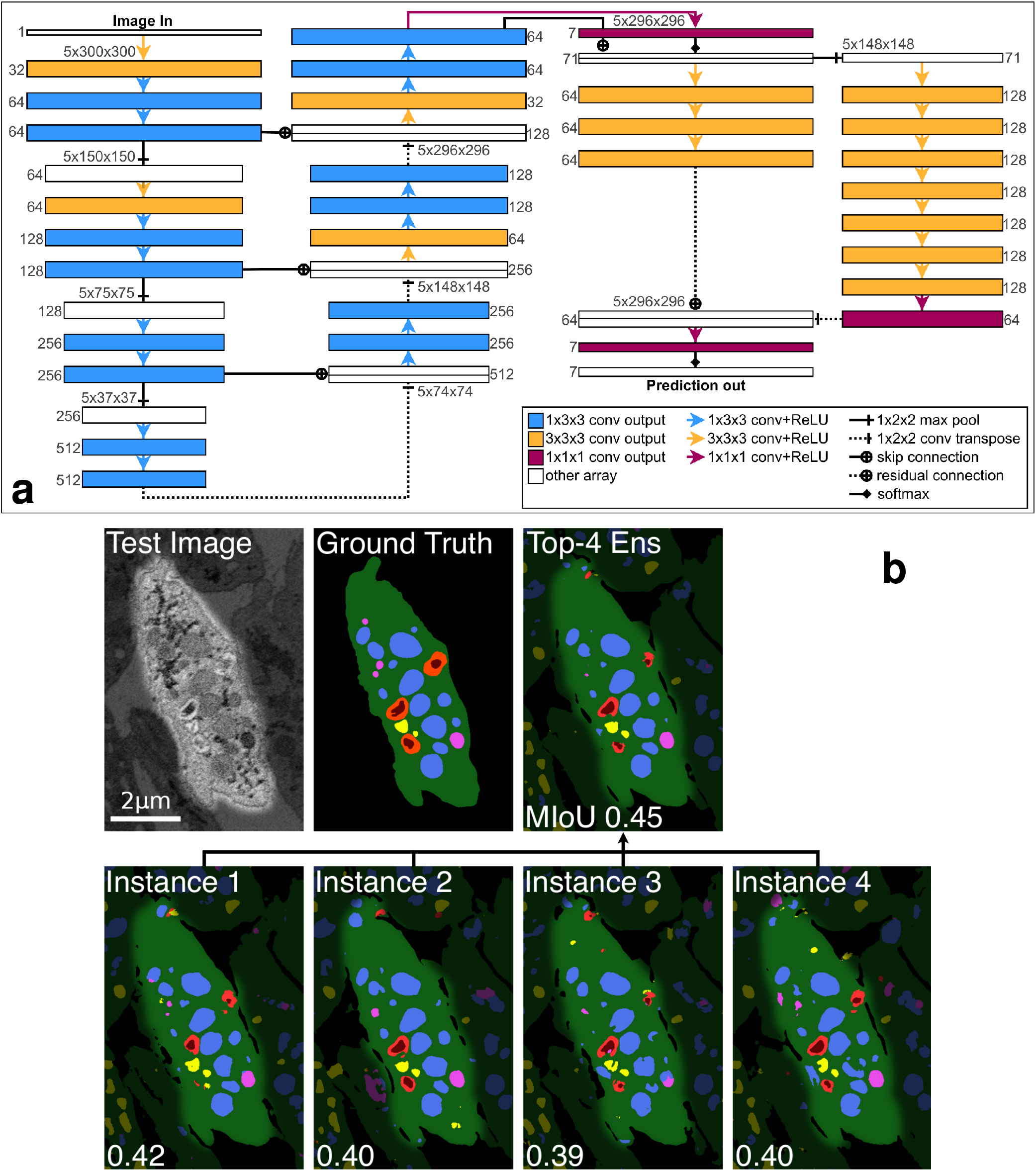
Methods. (a) Diagram of the 2D-3D+3×3×3 network architecture, the best design tested in this paper. Number triplets along box tops are layer spatial sizes. Numbers along box sides are layer convolution kernel counts. (b) Illustration of initialization-dependent performance of trained segmentation networks, and exploiting it for ensembling. An image of the test cell and ground truth labels are compared with segmentations of the best 4 trained 22D-3D+3×3×3 network instances and an ensemble formed from them. The ensemble improves MIoU^(*org*)^ by 7.1% over the best single network.

For the main experiment of this study, we train baseline architectures from the literature, our new architecture, and ablations of the new architecture on a dense cellular segmentation task by supervised learning from the training dataset. We compare single-network and ensemble segmentation performance on the evaluation and test datasets. We conclude that our algorithm outperforms baselines, the differentiating features of our final best architecture are responsible for the performance differences, and that multi-instance ensembles significantly improve performance over single networks. The results of this experiment are shown in Figure 1. We consider test performance to be the best indicator of an algorithm’s performance as it shows its ability to generalize across different samples. Figure 1 row 2 compares visualizations of the best 3D segmentation algorithms with ground-truth labels and image data for the test dataset. Figure 1 row 4 highlights the most notable performance results, and more performance statistics can be found in Table S1.

We also compare our best algorithm against the segmentations of scientific image annotators, Annotator 1 and Annotator 2, who are laboratory staff trained on annotation tasks but are not biological domain experts. These initial segmentations are currently the first step in producing high-quality dense cellular segmentations, and even before any corrections they require 1-2 work days per cell to create. Results are displayed in Figure 1 row 3, with further details in Figures S4 and S5. Annotator 1, Annotator 2, and our algorithm each labeled the annotator comparison region from the Subject2 platelet sample. We calculated MIoU^(*org*)^ scores pairwise from the three segmentations: 0.571 for Annotator 1 vs. Annotator 2, 0.497 for Annotator 1 vs. Algorithm, and 0.483 for Annotator 2 vs. Algorithm. The confusion matrices in Figure S5 further break down results by organelle class. The statistics indicate that our algorithm disagrees more with either annotator than the annotators do with each other, but none of the labels are consistent everywhere, reflecting the difficulty of dense cellular segmentation even for humans.

We are also interested in understanding how even imperfect segmentations may be useful for downstream analysis tasks. To this end, we computed organelle volume fractions for each organelle within the cell in the annotator comparison dataset. The cell volume fraction of an organelle is equal to the summed voxels of all organelles in a cell, divided by the cell’s volume. Biologists can correlate this information with other cell features to better understand variations in the makeup of cellular structures across large samples. The results in Figure 1 row 4 show that our algorithm tended to underestimate volume fractions relative to the two annotators, but the difference between the algorithm and Annotator 1 is smaller than the difference between Annotator 1 and Annotator 2. The best 3D algorithm improves considerably over the best 2D algorithm. All algorithms detect small regions ignored by humans, but simple postprocessing with small region removal fails to significantly improve quality metrics.

In conclusion, we have argued here that dense semantic labeling of 3D EM images for biomedicine is an image analysis method with transformative potential for structural biology. We demonstrated that while challenges exist for both human and algorithmic labelers, automated methods are approaching the performance of trained humans, and may be integrated into annotation software for greatly enhancing the productivity of humans segmenting large datasets. Without question, challenges remain for creating algorithms that are robust to the many types of variation present across research applications. However, SBF-SEM analysis problems are a fertile early ground for this computer vision research, as their large dataset sizes make the entire train-test-deploy cycle of supervised learning viable for accelerating analysis of even individual samples. We believe that the image in Figure 1(a-b) showcases this best - after manually segmenting less than 1% of the Subject 1 dataset, we were able to train a segmentation algorithm that produces a high-quality segmentation of the full dataset, a feat that would be impossible with anything short of an army of human annotators. While gains in accuracy will be realized with future developments, the procedure of training neural network ensembles on a manually annotated portion of a large SBF-SEM dataset is already becoming viable for making dense cellular segmentation a reality.

## Online content

Methods, supplementary materials, source data, code, and reproducible examples are available online at https://leapmanlab.github.io/dense-cell.

## Acknowledgments

This work was supported by the Intramural Research Program of the National Institute of Biomedical Imaging and Bioengineering, National Institutes of Health. We thank Prof. Brian Storrie and Dr. Irina Pokrovskaya, Department of Physiology and Biophysics, University of Arkansas for Medical Sciences, Little Rock, AR, for providing the embedded blocks of human blood platelets used in this study, and supported by NIH grant R01 HL119393. This work also used the computational resources of the NIH HPC Biowulf cluster. (https://hpc.nih.gov)

## Author Contributions

MDG and RDL conceived the project. BS and IDA conceived the data collection protocol and physically prepared the imaging samples. MDG, ZASE, and ABA designed and wrote the software. MDG, ZASE, and MAA performed experiments and processed and analyzed data. RDL and MDG coordinated the project. MDG wrote the paper.

## Methods

SBF-SEM image volumes were obtained from identically-prepared platelet samples from two humans. Lab members manually segmented portions of each volume into seven classes to analyze the structure of the platelets. The labels were used for the supervised training of candidate network architectures, as well as baseline comparisons. We trained multiple instances of candidate architectures, each with different random initializations. The best-performing instances were ensembled together to produce the final segmentation algorithms used in this paper.

### Data collection

This study used datasets prepared from two human platelet samples as part of a collaborative effort between the National Institute of Biomedical Imaging and Bioengineering (NIBIB), NIH and the University of Arkansas for Medical Sciences. The platelet samples were imaged using a Zeiss Sigma 3View SBF-SEM. The Subject 1 dataset is a 250 × 2000 × 2000 voxel image with a lateral resolution of 10nm and an axial resolution of 50nm, from a sample volume with dimensions 12.5 × 20 × 20*μ*m^3^. The Subject 2 dataset is a 239 × 2000 × 2000 voxel image produced by the same imaging protocol, with the same resolution and coming from a sample volume with the same dimensions. We assembled labeled datasets from manually-segmented regions of the platelet datasets. Lab members created initial manual segmentations using Amira^15^. For the training, evaluation, and test dataset, segmentations were reviewed by subject experts and corrected. The training image was a 50 × 800 × 800 subvolume of the Subject 1 dataset spanning the region 81 ≤ *z* ≤ 130, 1073 ≤ *y* ≤ 1872, 620 ≤ *x* ≤ 1419 in 0-indexed notation. The evaluation image was a 24 × 800 × 800 subvolume of the Subject 1 dataset spanning the region 100 ≤ *z* ≤ 123, 200 ≤ *y* ≤ 999, 620 ≤ *x* ≤ 1419. The test image was a 121 × 609 × 400 subvolume of the Subject 2 dataset spanning the region 0 ≤ *z* ≤ 120,460 ≤ *y* ≤ 1068, 308 ≤ *x* ≤ 707. The annotator comparison image was a 110 × 602 × 509 subvolume of the Subject 2 dataset spanning the region 116 ≤ *z* ≤ 225, 638 ≤ *y* ≤ 1239, 966 ≤ *x* ≤ 1474. The training and evaluation labels covered the entirety of their respective images, while the test and annotator comparison labels covered a single cell contained within their image volumes. The labeling schema divides image content into seven classes: `background` (0), `cell` (1), `mitochondrion` (2), `canalicular channel` (3), `alpha granule` (4), `dense granule` (5), and `dense granule core` (6).

### Neural architectures and ensembling

The Subject 1 and Subject 2 datasets were binned by 2 in *x* and *y*, and aligned. For each of the training, evaluation, and testing procedures, the respective image subvolumes were normalized to have mean 0 and standard deviation 1 before further processing.

The highest-performing network architecture in this paper, 2D-3D+3×3×3, is a composition of a 2D U-net-style encoder-decoder and 3D convolutional spatial pyramid, with additional 3×3×3 convolutions at the beginning of convolution blocks in the encoder-decoder. All convolutions are zero-padded to preserve array shape throughout the network, allowing deep architectures to operate on data windows with small **z**-dimension. A ReLU activation follows each convolution. All convolution and transposed convolutions use bias terms. The architecture is fully specified as a diagram in Figure 2. Additionally, several baseline comparison networks and three 2D-3D+3×3×3 ablation networks were also tested in this paper and are described in the Validation and Performance Metrics section. To build a 2D-3D network, one can adapt a 2D U-net-style encoder-decoder module to work on 3D data by recasting 2D 3×3 convolutions as 1×3×3 convolutions, and 2D 2×2 max-pooling and transposed convolution layers as 1×2×2 equivalents. In this way, a 3D input volume can be processed as a sequence of independent 2D regions in a single computation graph, and the 2D and 3D modules can be jointly trained end-to-end. Intermediate 2D class predictions 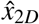 are formed from the 2D module output, and the 2D output and class predictions are concatenated along the feature axis to form an input to a 3D spatial pyramid module. The 3D module applies a 1×2×2 max pool to its input to form a two-level spatial pyramid with scales 0 (input) and 1 (pooled). The pyramid elements separately pass through convolution blocks, and the scale 1 block output is upsampled and added to the scale 0 block output with a residual connection to form the module output. 3D class predictions 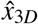 are formed from the 3D module output, and the final segmentation output 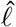 of the algorithm is a voxelwise argmax of the 3D class predictions. To build a 2D-3D+3×3×3 network, we inserted 3×3×3 convolution layers at the beginning of the first two convolution blocks in the 2D encoder and the last two convolution blocks in the 2D decoder.

Given a collection of networks’ 3D class predictions, one can form an ensemble prediction by computing a voxelwise average of the predictions and computing a segmentation from that. Ensembling high-quality but non-identical predictions can produce better predictions^16^, and there is reason to think that more sophisticated ensembles could be constructed from collections of diverse neural architectures^17^, but in this paper we use a simple source of differing predictions to boost performance: ensembles of identical architectures trained from different random initializations. The sources of randomness in the training procedure are examined more thoroughly in the Validation and Performance Metrics section, but in our experiments this variation produced a small number of high-performing network instances per architecture with partially-uncorrelated errors.

### Network training

We consider a network predicting classes *C =* {0,…,6} for each voxel in a shape-(*o_z_*,*o_x_*,*o_y_*) data window Ω containing *N* = *o_z_o_x_o_y_* voxels 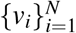. The ground-truth segmentation of this region is a shape-(*o_z_*, *o_x_*, *o_y_*) array *ℓ* such that *ℓ*(*v*) ∈ *C* is the ground-truth label for voxel *v*. A network output prediction is a shape-(7,*o_z_,o_x_,o_y_*) array 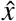 such that 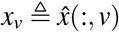 is a probability distribution over possible class labels for voxel *v*. The corresponding segmentation 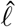 is the per-voxel arg max of 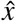. Inversely, from *ℓ* one may construct a shape-(7, *o_z_, o_x_, o_y_*) per-voxel probability distribution *x* such that *x_v_*(*i*) = 1 if *i* = *ℓ*(*v*) and 0 if not, which is useful during training.

We trained our networks as a series of experiments, with each experiment training and evaluating 1 or more instances of a fixed network architecture. Instances within an experiment varied only in the random number generator (RNG) seed used to control trainable variable initialization and training data presentation order. In addition to the main 2D-3D+3×3×3 architecture, there were three ablation experiments - No 3×3×3 Convs, No Multi-Loss, No 3D Pyramid - and five baseline experiments - Original U-Net, 3D U-Net Thin, 3D U-Net Thick, Deeplab + DRN, and Deeplab + ResNet101. Instances were trained and ranked by evaluation dataset MIoU. Experiments tracked evaluation MIoU for each instance at each evaluation point throughout training, and saved the final weight checkpoint as well as the checkpoint with highest evaluation MIoU. In this work we report evaluation MIoU checkpoints for each instance. The 2D-3D+3×3×3 experiment and its ablations trained 26 instances for 40 epochs with minibatch size 1 (33k steps). The Original U-Net experiment trained 500 instances for 100 epochs with minibatch size 1 (180k steps). The 3D U-Net Thin experiment trained 26 instances for 100 epochs with minibatch size 1 (29k steps), and the 3D U-Net Thick experiment trained 26 instances for 100 epochs with minibatch size 1 (30k steps). The Deeplab + DRN and Deeplab + ResNet101 experiments trained 1 instance each for 200 epochs with minibatch size 4 (360k steps). Due to poor performance and slow training times of the Deeplab models, we deemed it unnecessary to train further instances. Networks were trained on NVIDIA GTX 1080 and NVIDIA Tesla P100 GPUs.

This subsection details the training of the 2D-3D+3×3×3 network. Baseline and ablation networks were trained identically except as noted in Validation and Performance Metrics. All trainable variables were initialized from Xavier uniform distributions. Each instance was trained for 40 epochs on shape-(5,300,300) windows extracted from the training volume, and output a shape-(7,5,296,296) class prediction array. The number of windows in each epoch was determined by a window spacing parameter which determined the distance along each axis between the top-back-left corners of each window, here (2,100,100), resulting in 828 windows per epoch. An early stopping criterion halted the training of any network that failed to reach an MIoU of 0.3 after 10 epochs.

Networks were trained using a regularized, weighted sum of cross-entropy functions. The network has a set Θ trainable variables divided into four subsets: Θ_2*D*_ for variables in the 2D encoder-decoder module, Θ_3*D*_ for variables in the 3D spatial pyramid module, the single 1×1×1 convolution variable {*θ_2DP_*} which produces intermediate 2D class predictions 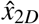 from the encoder-decoder’s 64 output features, and the single 1×1×1 convolution variable {*θ_3DP_*} which produces the final 3D class predictions 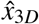 from the spatial pyramid’s 64 output features. Predictions are compared against ground-truth labels as

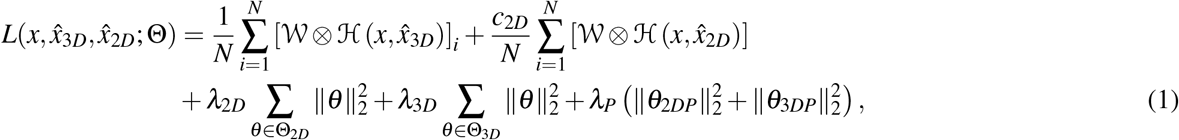

where λ_2*D*_ = 1 × 10^−4.7^ and λ_3*D*_ = 1 × 10^−5^ are **L**^2^ regularization hyperparameters for the variables in Θ_2*D*_ and Θ_3*D*_, *λ_P_* = 1 × 10^−9^ is an *L^2^* regularization hyperparameter for the predictor variables θ_2*DP*_ and θ_3*DP*_, and *c*_2*D*_ = 0.33 is a constant that weights the importance of the intermediate 2D class predictions in the loss function. 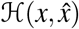 is the voxelwise cross-entropy function, i.e.,

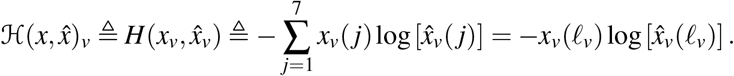

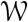 is a shape-(5,296,296) array of weights; its Kronecker product with 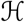 produces a relative weighting of the cross-entropy error per voxel. This weighting strategy is based generally on the approach in (Ronneberger et al., 2015)^4^:

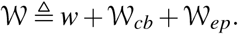

The initial *w* = 0.01 is a constant that sets a floor for the minimum weight value, 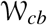 is a class-balancing term such that 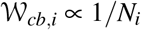, where *N_i_* is the number of occurrences in the training data of *ℓ_i_*, rescaled so that max 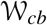 = 1. 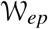 is an edge-preserving term that upweights voxels near boundaries between image objects and within small 2D cross-sections. In (Ronneberger et al., 2015) this is computed using morphological operations. We used a sum of scaled, thresholded diffusion operations to approximate this strategy in a manner that requires no morphological information. 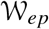 is built up as a rectified sum of four terms:

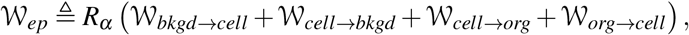

where 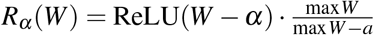. For each term, we choose two disjoint subsets *C_source_* and *C_target_* of the *C*. Let *ℓ_source_* be the binary image such that *ℓ_source_*(*v*) = 1 if ℓ(*v*) ∈ *C_source_* and *ℓ_source_*(*v*) = 0 otherwise. Define

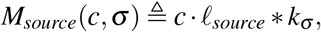

where * denotes convolution and *k_σ_* is a Gaussian diffusion kernel with standard deviation σ. Then, 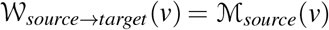 if ℓ(*v*) ∈ *C_target_*, and is 0 otherwise. The terms in the 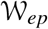 array used in this paper were computed using class subsets *bkgd* = {0}, *cell* = {1}, and *org* = {2,3,4,5,6}, *α* = 0.25, *c* = 0.882, and *σ* = 6. The error weighting array used in this paper and the code used to generate it are available with the rest of the platelet dataset at leapmanlab.github.io/dense-cell. See Figure S7 for a visualization of the error weighting array. 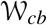 is calculated all at once across the entire 3D training volume, while 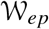 is calculated independently per each 2D *z*-slice of the training volume.

We employed data augmentation to partially compensate for the limited available training data. Augmentations were random reflections along each axis, random shifts in brightness (±12%) and contrast (±20%), and elastic deformation as in (Ronneberger et al., 2015). For elastic deformation, each 800×800 *x – y* plane in the shape- (50, 800, 800) training data and label arrays was displaced according to a shape-(800, 800, 2) array of 2D random pixel displacement vectors, generated by bilinearly upsampling a shape-(20,20,2) array of iid Gaussian random variables with mean 20 and standard deviation 0.6. During each epoch of training, a single displacement map was created and applied to the entire training volume before creating the epoch’s batch of input and output windows. Training used the ADAM optimizer with learning rate 1 × 10^−3^, *β*_1_ = 1 − 1 × 10^−1.5^, *β*_2_ = 1 − 1 × 10^−2.1^, and *ε* = 1 × 10^−7^. Training also used learning rate decay with an exponential decay rate of 0.75 every 1 × 10^3.4^ training iterations.

### Validation and performance metrics

The performance metric used in this work is mean intersection-over-union (MIoU) between ground-truth image segmentation *ℓ*’s 7 labeled sets {*L_j_* = *v* ∈ Ω | *ℓ*(*v*) = *j*}_*j*∈*C*_ and predicted segmentation’s 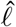 labeled sets 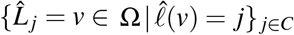. Given two sets *A* and *B*,

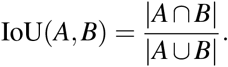

Then for segmentations *ℓ* and 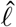 with their corresponding labeled sets,

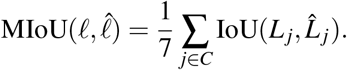

More generally, for a subset of labels *D* ⊆ *C*, one can compute the MIoU over *D*, or MIoU^(*D*)^, as

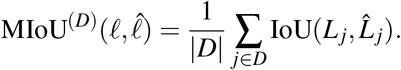

Here we are concerned with MIoUs over two sets of labels: MIoU^(*all*)^ over the set of all 7 class labels, and MIoU^(*org*)^ over the set of 5 organelle labels 2-7. Our network validation metrics were MIoU^(*all*)^ and MIoU^(*org*)^ on the evaluation dataset, and MIoU^(*org*)^ on the test dataset. Test data uses MIoU^(*org*)^ because the labeled region is a single cell among several unlabeled ones, and restricting validation to the labeled region invalidates MIoU stats for the background and cell classes (0 and 1). We include evaluation MIoU^(*org*)^ to quantify how performance drops between a region taken from the physical sample used to generate the training data, and a new physical sample of the same tissue system. Using this procedure, the performance of the 2D-3D+3×3×3 network was compared against three ablations and five baseline networks. The three ablations each tested one of three features that distinguish the 2D-3D+3×3×3 network in this paper from similar baselines. The first, 2D+3×3×3 No 3×3×3 Convs, replaces the 3×3×3 convolutions in the net’s encoder-decoder module with 1×3×3 convolutions that are otherwise identical. With this ablation, the network’s encoder-decoder loses any fully-3D layers. The second, 2D+3×3×3 No Multi-Loss, modifies the loss function in Equation (1) by removing the term involving *x*2*D* but otherwise leaving the architecture and training procedure unchanged. This ablation tests whether it is important to have auxiliary accuracy loss terms during training. The third ablation, 2D-3D+3×3×3 No 3D Pyramid, removes the 3D spatial pyramid module and 3D class predictor module from the network architecture, so that 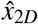 is the network’s output. Correspondingly, the loss term involving 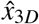 is removed from Equation (1).

We implemented five baseline networks by adapting common models in the literature to our platelet segmentation problem. Three of these were 2D - The original U-Net^4^ as well as two Deeplab variants^18,19^ using a deep residual network (DRN) backbone and a ResNet101 backbone^20^, minimally modified to output 7 class predictions. The original U-Net used (572,572) input windows and (388,388) output windows, while the Deeplab variants used (572,572) input and output windows. The two 3D networks were fully-3D U-Net variants adapted on the 3D U-Net in (Çiçek et al., 2016)^6^ - 3D U-Net Thin and 3D U-Net Thick. The variants used same-padding, had three convolutions per convolution block, and two pooling operations in the encoder for convolution blocks at three spatial scales. The 3D U-Net Thin network used (5,300,300) input windows and (5,296,296) output windows, and pooling and upsampling operations did not affect the *z* spatial axis. The 3D U-Net Thick network used (16,180,180) input windows and (16,180,180) output windows, and pooled and upsampled along all three spatial axes.

To determine whether one architecture is superior to another, trained instances are compared with each other. However, sources of randomness in the training process induce a distribution of final performance metric scores across trained instances of an architecture, so that a single sample per architecture may be insufficient to determine which is better. While expensive, a collection of instances can be trained and evaluated to empirically approximate the performance distribution for each architecture. In this way, better inferences may be made about architecture design choices. Figure S6 shows the empirical performance distributions for the 26 trials of the 2D-3D+3×3×3 architecture and its three ablations, as well as the 26 trials of the 3D U-Net and 500 trials of the 2D Original U-Net. A final point of comparison was drawn between the top algorithm’s performance and the initial work of laboratory scientific image annotators, Annotator 1 and Annotator 2. The three sources each labeled the annotator comparison region from the Subject 2 platelet sample. Pairwise MIoU^(*org*)^ scores and organelle confusion matrices were calculated to compare the level of disagreement between two human labelings and between humans and the algorithm. We also computed organelle volume fractions for each segmentation to compare performance in segmentation applications to downstream analysis tasks. The cell volume fraction of an organelle is equal to the summed voxels of all organelles in a cell, divided by the cell’s volume. To compute this quantity for each organelle, the number of voxels for each organelle label is divided by the number of voxels in the cell. For the algorithmic result, since the semantic segmentation map does not distinguish between separate cells in the field of view, a mask for the single annotator comparison dataset cell was approximated as all non-background-labeled voxels in a small region around the Annotator 1 cell mask.

In addition to the multiclass baselines, we chose to also evaluate a CDeep3M plug-and-play system^12^ that can be spun up on Amazon Web Services (AWS) for binary segmentation problems. We compared the full CDeep3M 3D network ensemble trained for 30000 iterations and all other default settings against our best 3D ensemble on a binarized version of the evaluation dataset. The ground truth labels and our ensemble’s output were binarized by mapping all labels greater than 0 (all cell content) to 1. The CDeep3M cell content predictions were detection probabilities between 0 and 1, and we computed the MIoU for CDeep3M predictions at 11 equispaced detection thresholds from between 0 and 1. Results did not change the final conclusions about our work, and so details are in Supplementary Materials.

### Supplementary Material

#### Additional segmentation visualizations

In addition to the renderings in the main paper, we produced 3D renderings of segmentation results for the evaluation dataset, as shown in Figure S2, showing results for all organelles together as well as separately for the ground-truth labels, as well as the best 2D and 3D segmentation algorithms. Similarly, Figure S4 shows renderings per each organelle class for Annotator 1, Annotator 2, and our best algorithm on the annotator comparison dataset. Finally, Figure S3 shows 2D images of segmentations for each of the 14 algorithms tested in this paper, which are also detailed in Table S1.

#### Ablation analysis and initialization-dependent performance

Our ablation analysis procedure, described in the **Validation and performance metrics** subsection of **Methods** confirms our conjectures about the importance of 3D context input to the network, and the importance of 3×3×3 convolutions over 1×3×3 convolutions for generalization performance. The latter do not capture correlations along the *z* spatial dimension, likely contributing to their poorer performance. Ablation analysis also indicates that removing either the multi-loss training setup or the 3D spatial pyramid module from the 2D-3D+3×3×3 architecture carries significant performance penalties. Removing either the 3×3×3 convolution layers or the 3D spatial pyramid on their own had a small effect on performance compared with removing the 2D loss term from the multi-loss objective function. Summary statistics demonstrating these results can be seen in Table S1(a), but these statistics only tell part of the story. Especially when effect sizes are small, looking at a single trained instance of each architecture may not be enough to determine relative performance between architecture candidates. To get a better idea of the effects of different architecture choices, we must deal with the initialization-dependent performance of these segmentation networks.

In Figure S6 we experiment with various weight initialization random seeds to determine the robustness of various models to the weight initialization scheme. In order to determine whether one architecture choice is superior to another, the outputs of different trained networks are compared with each other. However, sources of randomness in the training process (initialization of trainable weights from a Xavier uniform distribution, and the random presentation order of training data elements) induce a distribution of final performance metric scores. These scores are random variables, and a single sample per architecture may be insufficient to determine which is better. By empirically approximating the distribution for each architecture, better inferences may be made about architecture design choices. For this figure, multiple instances of the same architecture (26 for 2D-3D and fully-3D nets, 500 for the U-Net) were trained under identical conditions, varying only random number generation seeds. The resulting distributions support the conclusions that 2D-3D networks outperform their 2D and fully-3D counterparts, as well as the conclusions drawn from the ablation studies. They also reveal a curious phenomenon that may be a topic for future study - the seemingly bimodal performance of 2D-3D architectures, wherein some fraction of trained instances perform markedly worse than others with an apparent performance gap between peaks. Whether this is a real phenomenon or an artifact of having an insufficient number of samples could be determined with a followup study.

#### DeepVess baseline comparison

In addition to the baseline models discussed in the main text, we have also tried using the DeepVess model from (Haft-Javaherian et al., 2019)^8^ on our data. However, DeepVess performed poorly, and learned to assign a single class (background) to the entire output patch. We believe there may be two reasons behind DeepVess’ poor performance: (1) Unlike U-Net and Deeplab networks, the DeepVess network is designed with very small input patches in mind; small patches do not contain enough context for the network to distinguish between objects. (2) DeepVess’ last layer consists of a fully-connected operation with a single hidden layer containing 1024 neurons, therefore any attempt to input significantly larger patches would require increasing the number of neurons in the last layer, but fully-connected layers do not scale well and the network quickly outgrows GPU memory.

#### CDeep3M baseline comparison

A recent addition to the microscopy segmentation literature and software ecosystem is the CDeep3M tool from Haberl et al.^12^. In a similar vein to our work and others’, they use an ensemble of convolutional neural networks to perform binary segmentation tasks. This differs from the multiclass segmentation problems that we address, but their polished workflow makes it easy to replicate and train on new data. We therefore decided to evaluate CDeep3M on a comparable binary segmentation task with our data, wherein all non-`background` classes were grouped together into a single `cell` class. Using the AWS stack provided on the project GitHub page (https://github.com/CRBS/cdeep3m), we trained the networks used in their 3D segmentation ensemble for 30000 iterations on our training dataset, using all other default hyperparameters. Training took approximately 96 hours on an Amazon EC2 instance with an NVIDIA P100 GPU card.

After training completed, we ran the CDeep3M 3D ensemble’s prediction tool on our evaluation dataset, and compared it with a binarized version of our best algorithm’s segmentation of the evaluation dataset. The results can be seen in Figure S8. We binarized our algorithm’s segmentation the same way we binarized our ground truth labels, by mapping together all the non-`background` segmented classes. The CDeep3M algorithm, however, produces a single per-voxel probability map that indicates the probability each voxel belongs to a cell region. To compute a segmentation from the probability map, a cutoff threshold *t* must be specified - a segmentation with threshold *t* assigns the `cell` class to all voxels with probability greater than *t*, and `background` to all others. We computed MIoU scores for our lab’s (LCIMB) segmentation, as well as CDeep3M segmentations with thresholds in {0,0.1,0.2,0.3,0.4,0.5,0.6,0.7,0.8,0.9,1}. The 0.4 and 0.5 thresholds both produced the highest MIoU score - 0.935. In contrast, the LCIMB segmentation had an MIoU of 0.946. Both algorithms generally did a good job of detecting cell material, but the LCIMB segmentation did a much better job of preserving boundaries between adjacent cells.

#### Segmentation 3D rendering videos

In addition to the 3D rendering images of segmentations displayed in figures in this work, we produced videos showing rotations of the renderings.

#### Evaluation dataset

Ground truth: https://leapmanlab.github.io/dense-cell/vids/eval_gt.mp4

Best 3D ensemble: https://leapmanlab.github.io/dense-cell/vids/eval_e-3d.mp4

Best 2D ensemble: https://leapmanlab.github.io/dense-cell/vids/eval_e-2d.mp4

Best 3D network: https://leapmanlab.github.io/dense-cell/vids/eval_s-3d.mp4

Best 2D network: https://leapmanlab.github.io/dense-cell/vids/eval_s-2d.mp4

#### Test dataset

Ground truth: https://leapmanlab.github.io/dense-cell/vids/test_gt.mp4

Best 3D ensemble: https://leapmanlab.github.io/dense-cell/vids/test_e-3d.mp4

Best 2D ensemble: https://leapmanlab.github.io/dense-cell/vids/test_e-2d.mp4

Best 3D network: https://leapmanlab.github.io/dense-cell/vids/test_s-3d.mp4

Best 2D network: https://leapmanlab.github.io/dense-cell/vids/test_s-2d.mp4

#### Annotator comparison dataset

Annotator 1: https://leapmanlab.github.io/dense-cell/vids/ac_ann1.mp4

Annotator 2: https://leapmanlab.github.io/dense-cell/vids/ac_ann2.mp4

Best algorithm: https://leapmanlab.github.io/dense-cell/vids/ac_alg.mp4

#### Training demonstration videos

We trained a 2D-3D+3×3×3 network for 39744 iterations, recording the class prediction probability maps and segmentation that the network produced on the evaluation dataset every 92 iterations. We produced animations of the evolution of the network’s prediction capabilities to demonstrate the learning process.

#### Probability maps video

The first video shows the evolution of the six non-background probability maps predicted by the network over the course of training. Each probability map is color-coded based on the corresponding structure color in the segmentation color scheme used throughout this paper - dark green for `cell`, magenta for `mitochondrion`, dark blue for `alpha granule`, yellow for `canalicular channel`, bright red for `dense granule`, and dark red for `dense granule core`.

Link: https://leapmanlab.github.io/dense-cell/vids/train_prob-maps.mp4

#### Segmentation video

The second video shows the evolution of the segmentation produced by the network over the course of training. Link: https://leapmanlab.github.io/dense-cell/vids/train_seg.mp4

---

**Table S1.**
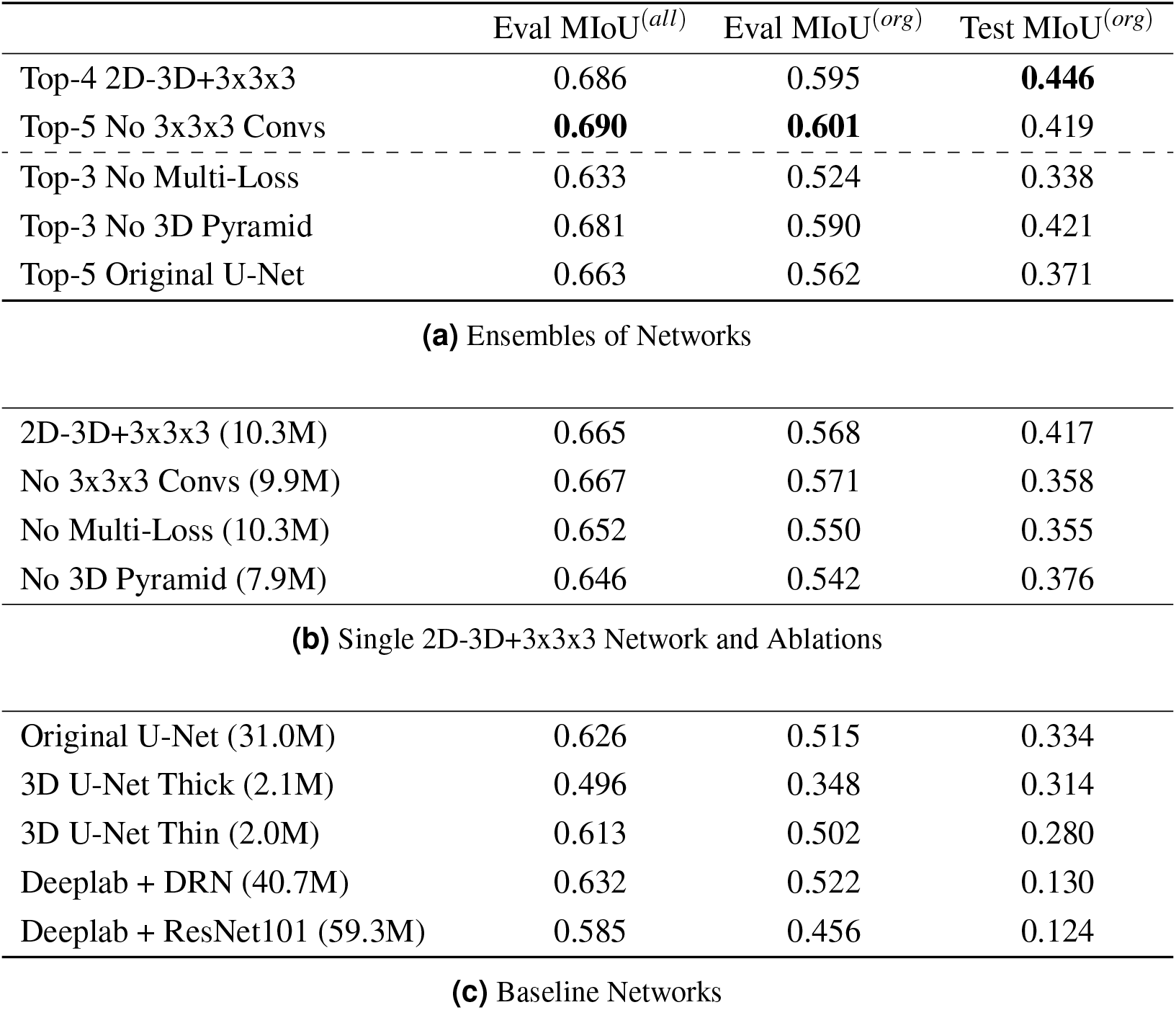
Comprehensive network performance statistics. Segmentation algorithm results summary showing mean intersection-over-union (MIoU) across all classes (MIoU^(*all*)^) on evaluation data and MIoU across organelle classes (MIoU^(*org*)^) on evaluation and test data. The Subject 2 dataset from which the test data is taken contains only a small number of labeled cells among unlabeled ones; we use MIoU^(*org*)^ to measure test performance since restricting the MIoU stat to labeled regions invalidates `background` and `cell` class statistics. (**a**) Results for the best ensemble from each architecture tested. A top-*k* ensemble averages the predictions of the best *k* trained networks as judged by MIoU^(*all*)^ on the evaluation dataset. (**b**) Results for the best single network from each architecture class. Trainable parameter counts are in parentheses. (**c**) Results from baseline comparison networks. Trainable parameter counts are in parentheses

**Figure S1.**
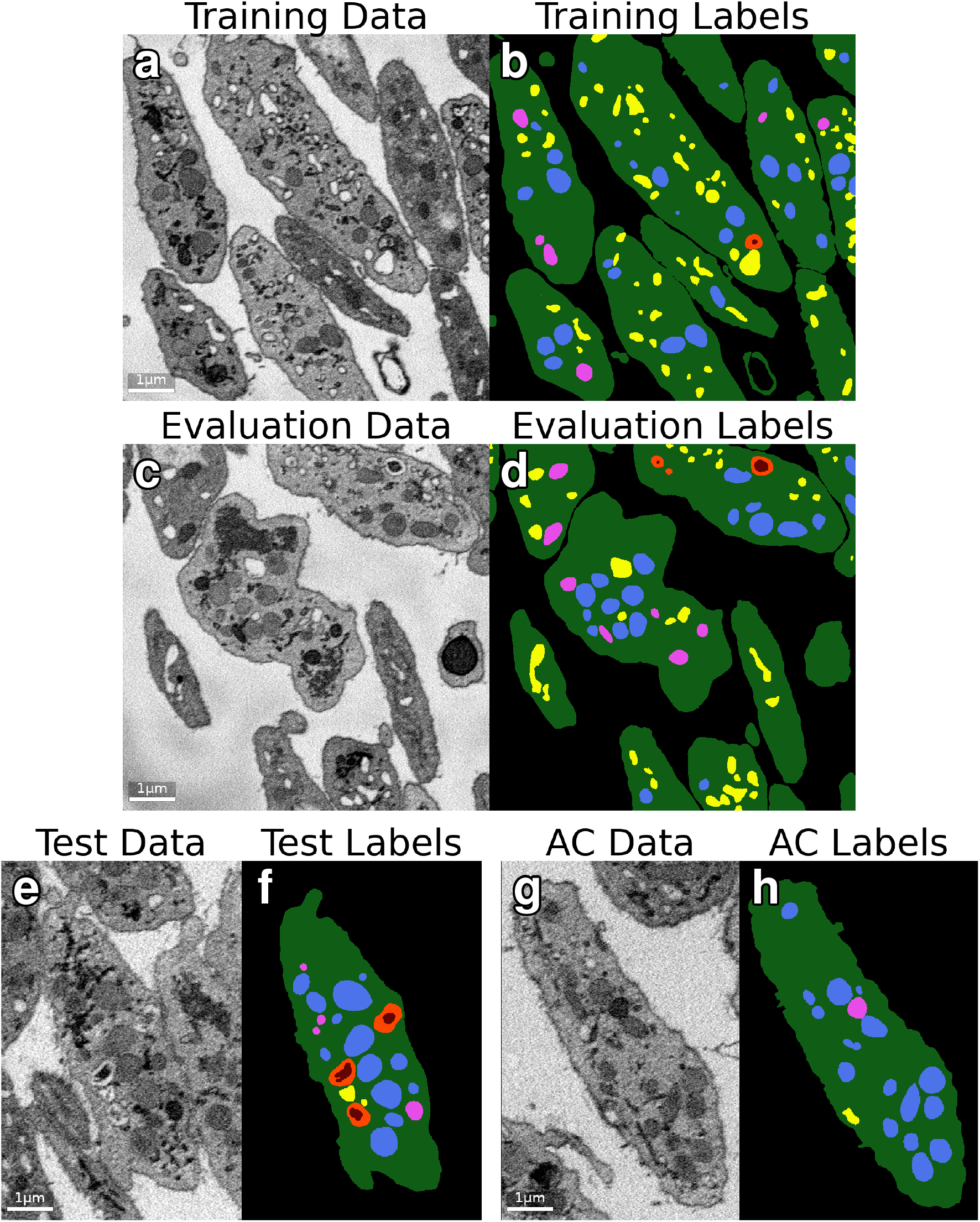
Dataset visualization. Sample orthoslices of the datasets used in this study. (**a-b**) Training image data and labels. (**c-d**) Evaluation image data and labels. (**e-f**) Test image data and labels. (**g-h**) Annotator comparison (AC) image data and labels.

**Figure S2.**
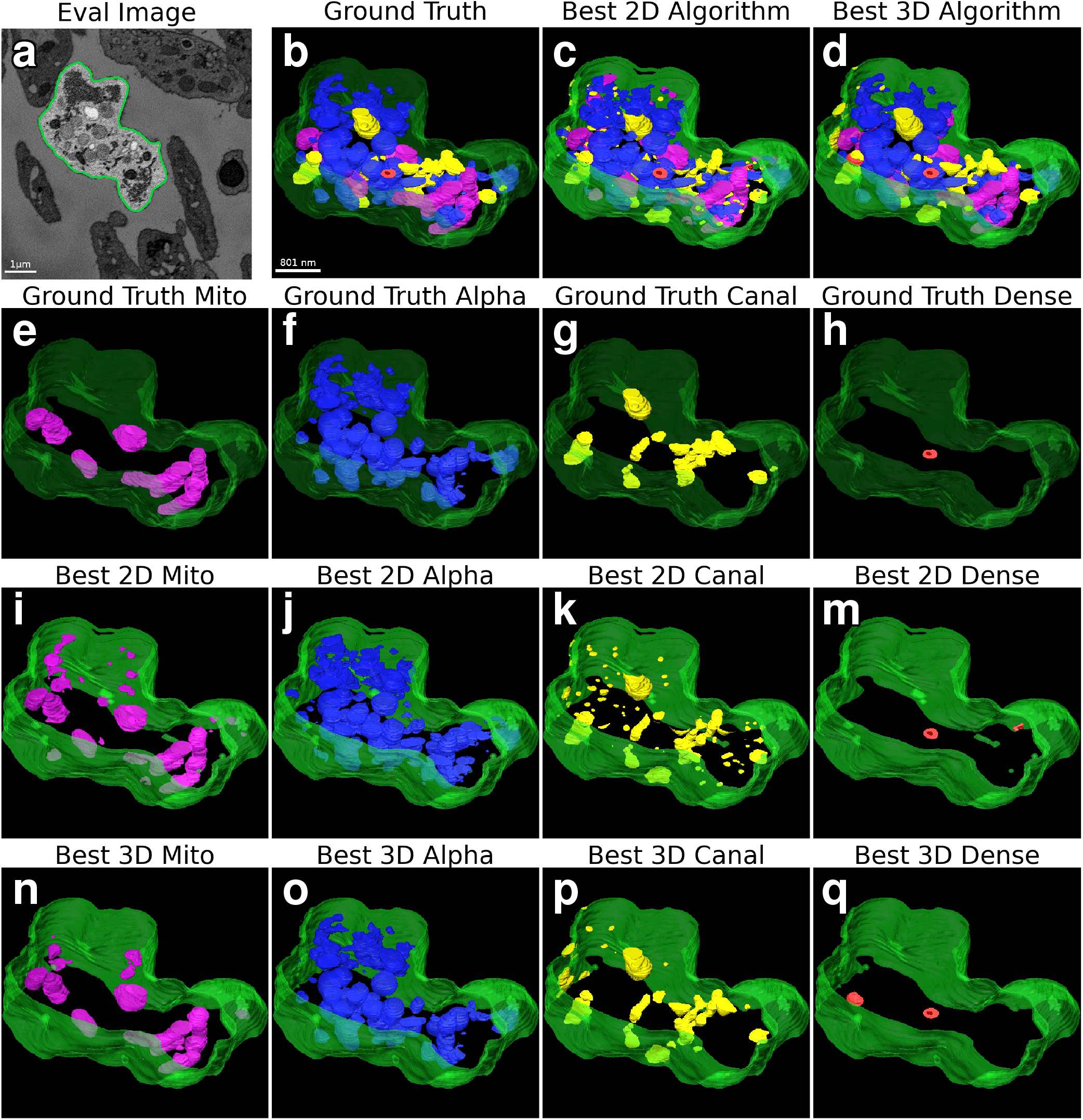
Evaluation dataset segmentation renderings. (**a**) Orthoslice of the evaluation dataset with rendered cell highlighted. (**b-d**) Ground truth and best 2D and 3D algorithm segmentations of the evaluation cell region showing all organelles. (**e-h**) Ground truth segmentations of individual organelles - from left to right: mitochondria (Mito), alpha granules (Alpha), canalicular channels (Canal), dense granules (Dense). (**i-m**) Best 2D algorithm segmentations of individual organelles. (**n-q**) Best 3D algorithm segmentations of individual organelles

**Figure S3.**
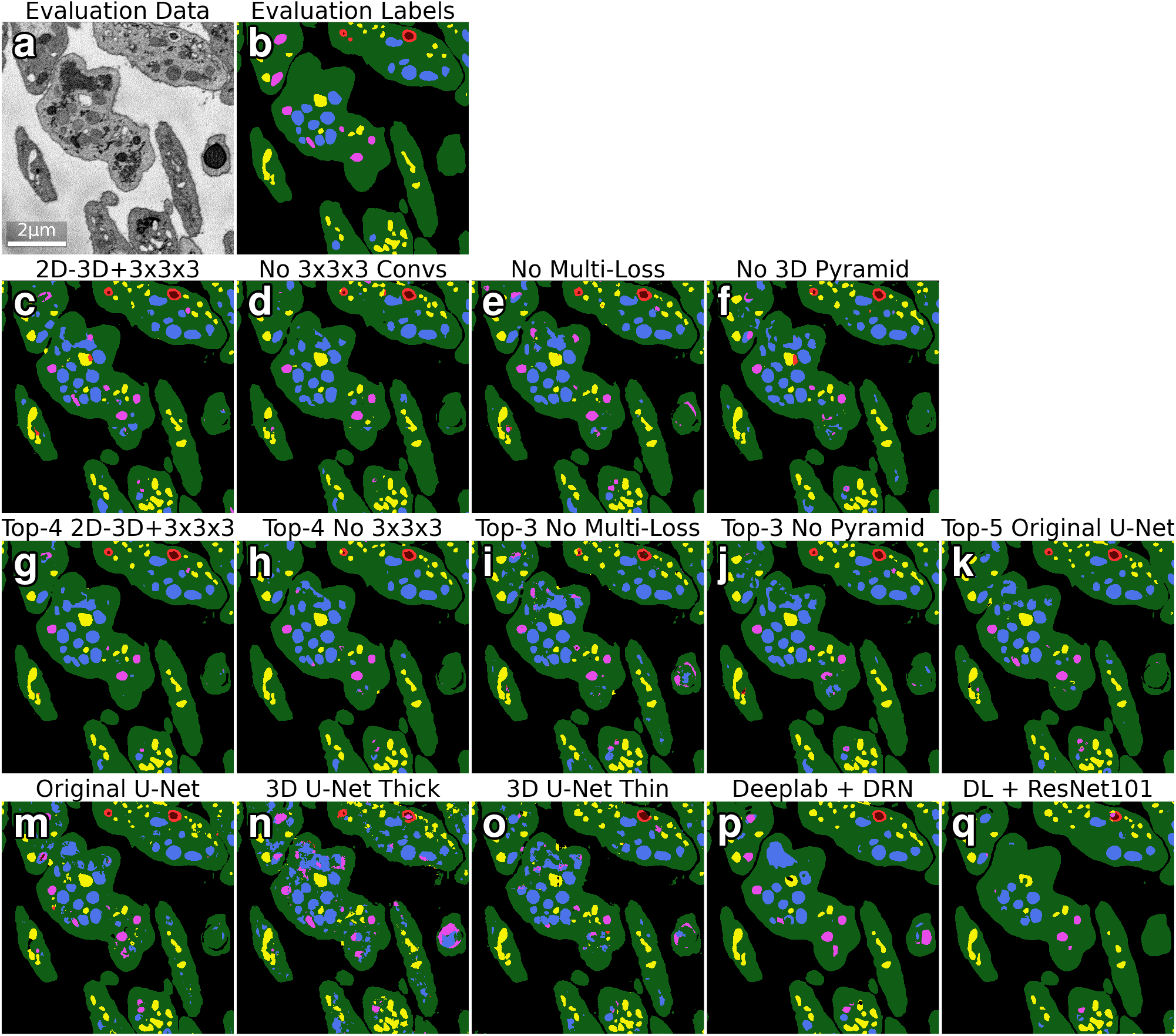
2D comparison of all algorithm results. This figure compares the results of all 14 segmentation algorithms tested in this paper with ground-truth labels for the *z* = 4 slice of the evaluation dataset. (**a-b**) Orthoslice of the evaluation image dataset and segmentation. (**c-f**) Segmentations from our new 2D-3D+3×3×3 network and its three ablations. (**g-k**) Segmentations from the five ensemble algorithms tested in this work. (**m-q**) Segmentations from the five baseline networks tested in this work.

**Figure S4.**
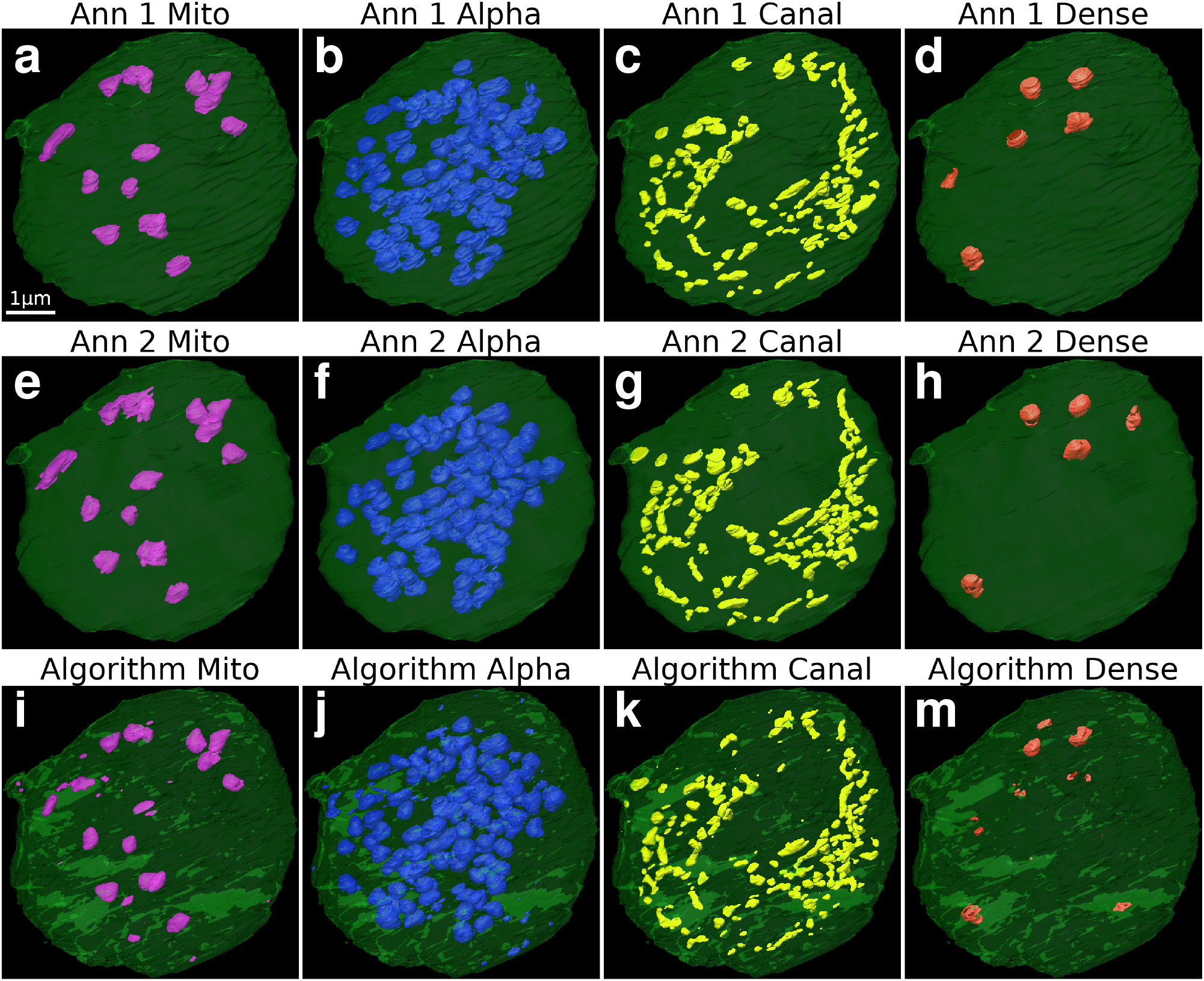
Annotator comparison segmentation renderings. This figure supplements row 4 of Figure 1 by showing renderings of individual organelle types - Mito for mitochondria, Alpha for alpha granules, Canal for canalicular channels, Dense for dense granules - from the Annotator 1, Annotator 2, and best Algorithm (Top-4 2D-3D+3×3×3) segmentations. (**a-d**) Annotator 1 (Ann 1) organelle segmentations. (**e-h**) Annotator 2 (Ann 2) organelle segmentations. (**i-m**) Algorithm organelle segmentations.

**Figure S5.**
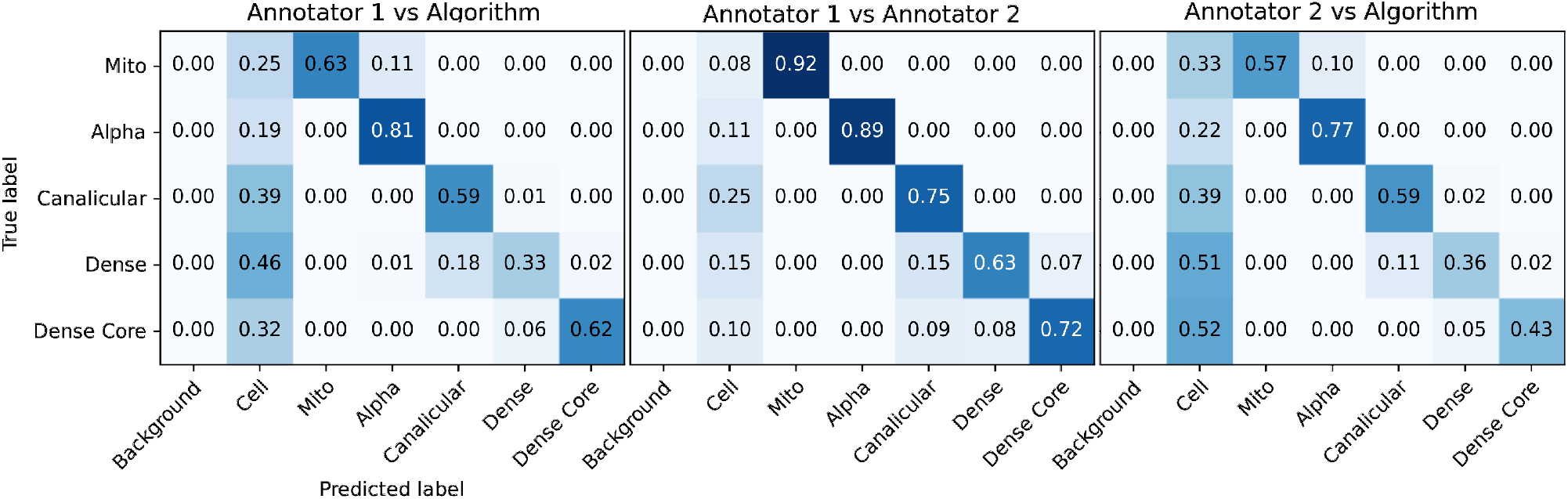
Annotator comparison confusion matrices. Confusion matrices comparing organelle labelings pairwise between the two annotators and our best algorithm. These give a more detailed performance breakdown of the MIoU^(*org*)^ scores obtained for each comparison: 0.497 for Annotator 1 vs Algorithm, 0.571 for Annotator 1 vs Annotator 2, and 0.483 for Annotator 2 vs Algorithm.

**Figure S6.**
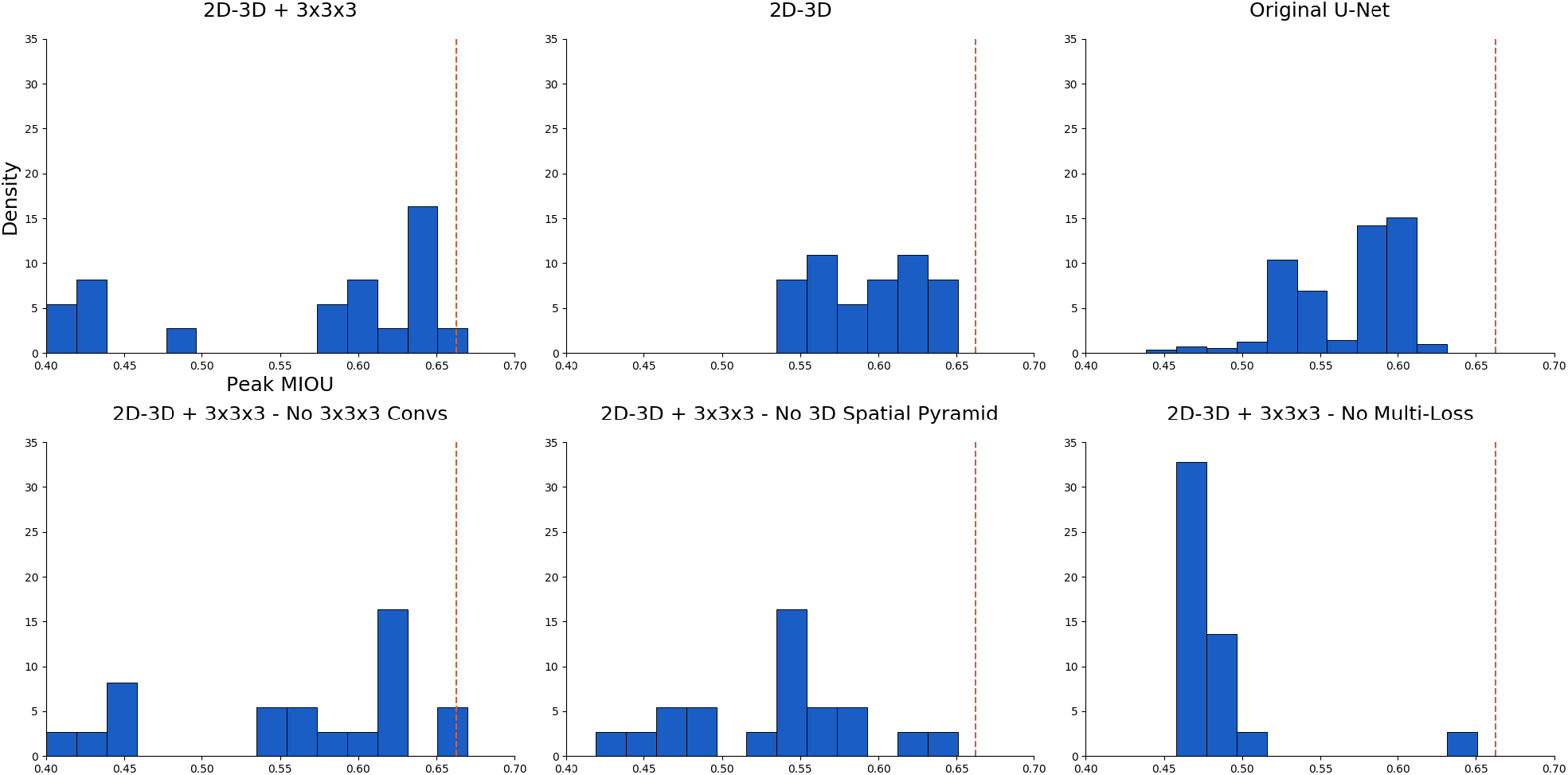
Making better architecture design decisions. This figure shows normalized histograms of peak MIoU^(*all*)^ on the evaluation dataset for each of the architectures examined in this paper. In order to determine whether one architecture choice is superior to another, the outputs of different trained networks are compared with each other. However, sources of randomness in the training process (initialization of trainable weights from a Xavier uniform distribution, and the random presentation order of training data elements) induce a distribution of final performance metric scores. These scores are random variables, and a single sample per architecture may be insufficient to determine which is better. By empirically approximating the distribution for each architecture, better inferences may be made about architecture design choices. For this figure, multiple instances of the same architecture (26 for 2D-3D nets, 500 for the U-Net) were trained under identical conditions, varying only random number generation seeds. The resulting distributions support the conclusions that 2D-3D networks outperform the 2D U-Net and that multi-loss training is necessary for 2D-3D architectures.

**Figure S7.**
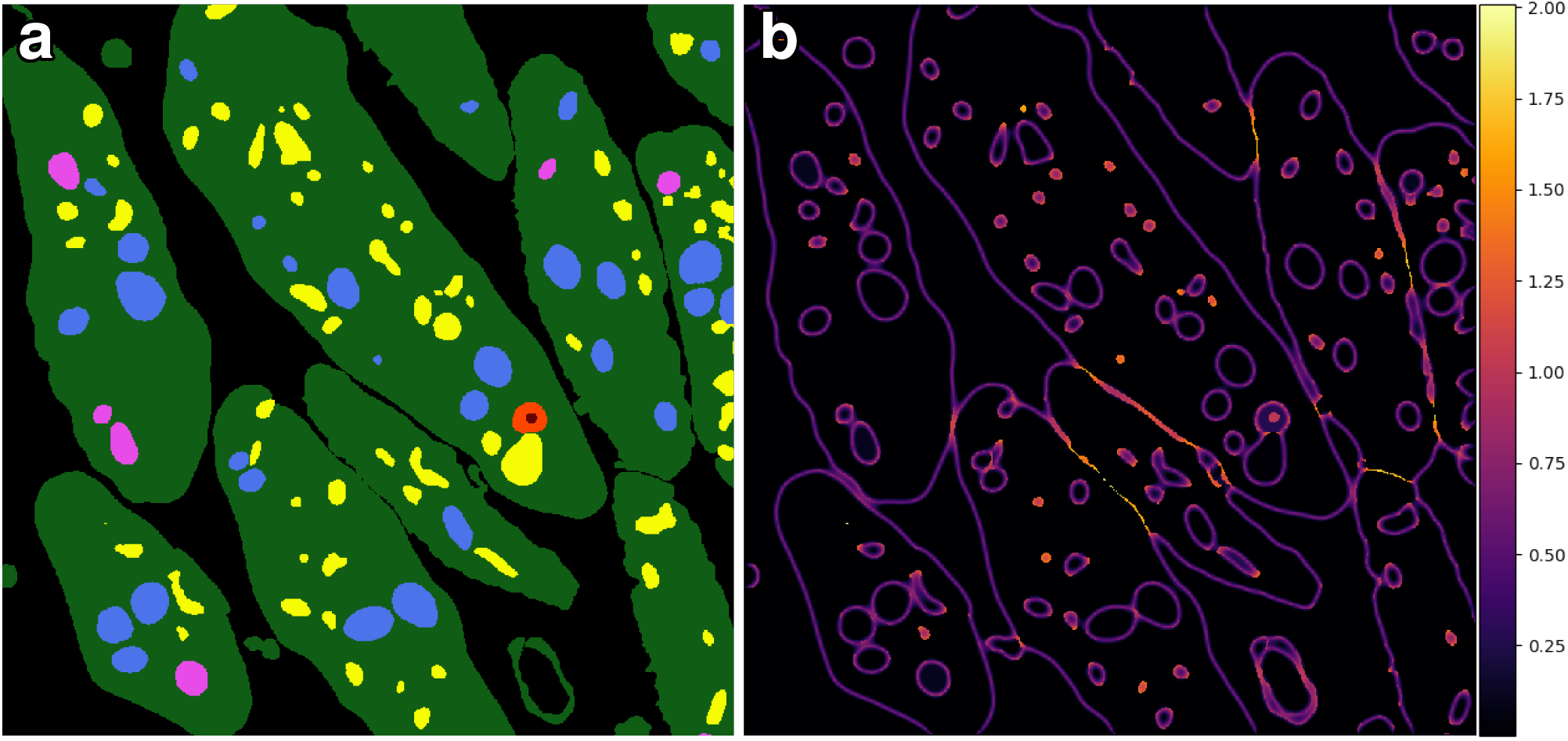
Error weighting array visualization. The error weighting 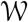 array is the sum of three terms 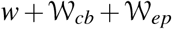, where *w* is a weight floor, 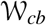 is a class frequency balancing array, and 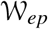 is an edge preserving array. 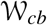 and 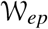 are computed from ground truth labels. (**a**) Example orthoslice of training dataset ground truth labels. (**b**) Corresponding orthoslice of the error weighting array.

**Figure S8.**
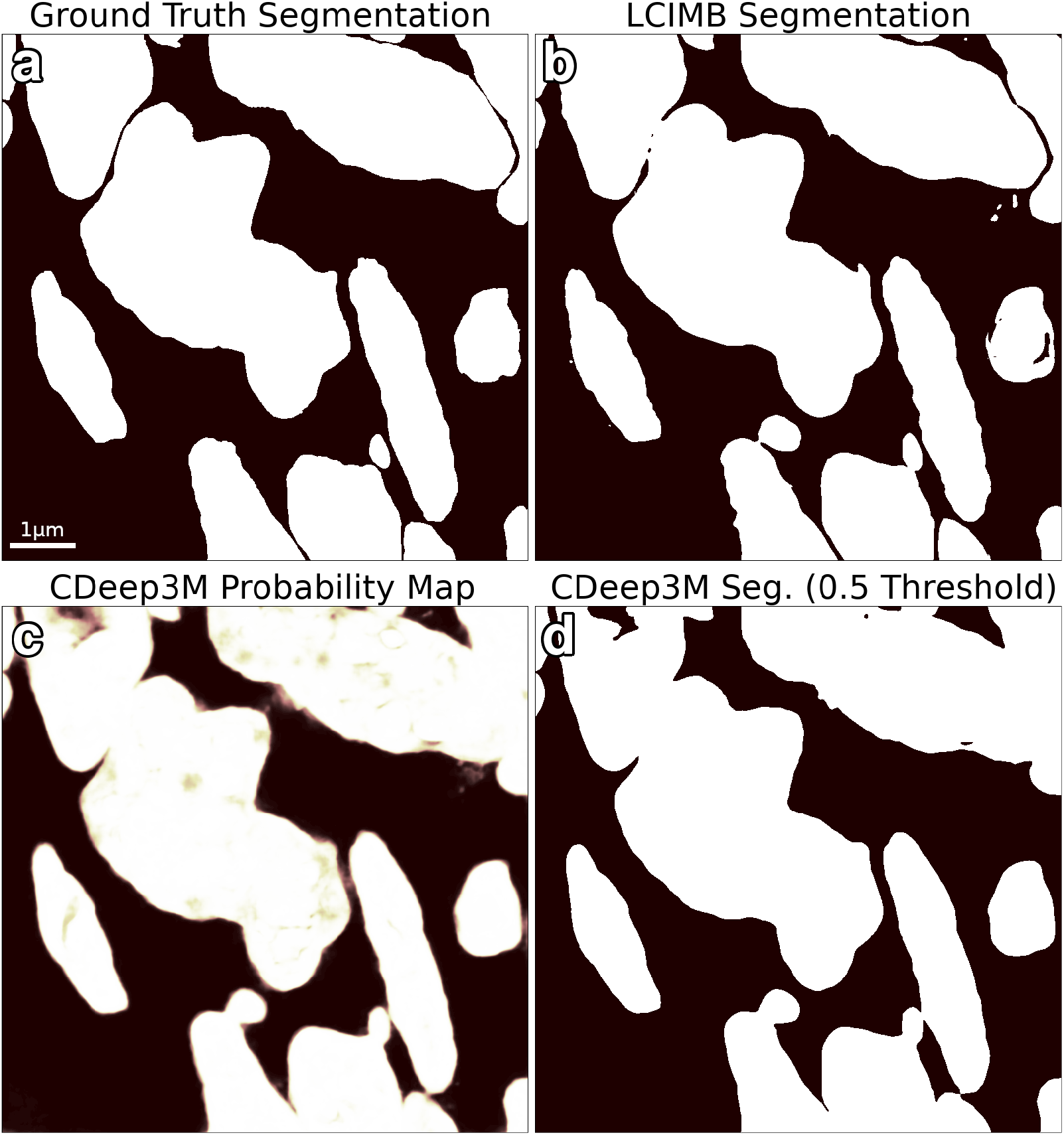
CDeep3M segmentation comparison. Comparison between the CDeep3M segmentation tool and our lab’s (LCIMB) best segmentation algorithm for a binary cell/non-cell segmentation problem on our evaluation dataset. (**a**) Orthoslice of the ground truth binary segmentation of the evaluation dataset. (**b**) Segmentation using our lab’s (LCIMB) best 3D ensemble. (**c**) Probability map produced by the CDeep3M ensemble after training on our data for 30000 iterations. The probability map is a per-voxel probability that the voxel belongs to a cell region, and it must be thresholded to produce a segmentation. (**d**) Segmentation from the CDeep3M ensemble with the best tested threshold of 0.5. This resulted in an MIoU of 0.935, compared to 0.946 for the LCIMB segmentation. In addition to a slight improvement in MIoU statistic, the LCIMB segmentation does a much better job of preserving boundaries between adjacent cells

